# A chemical space model for the exploration of eco-toxicological data

**DOI:** 10.1101/2025.09.18.677052

**Authors:** D Lopez-Rodriguez, G Guerrero-Limón, N Chèvre

## Abstract

With over 350000 chemicals and mixtures currently registered for production and use worldwide, up to 90% of authorized chemicals lack adequate toxicity data, leaving the majority of chemicals poorly characterized. International agencies urge scientists to develop screening methods to explore, identify, and predict chemical hazards, supporting the prioritization of chemical risk assessment. Here, Tree Manifold Approximation and Projection (TMAP) were applied, with the aim of reducing the dimensionality of large toxicological dataset, providing the foundations to data imputation methods allowing to get an understanding of chemical modes of action. Specifically, TMAP was implemented using MHFP6 fingerprints and the NORMAN SusDat database, containing over 100000 compounds. Physicochemical properties and CTD toxicogenomic data were retrieved and a graph-based spatial imputation function was generated to obtain insights into the potential ecotoxicity mechanisms of data poor chemicals. The relevance and meaningfulness of TMAP chemical space was explored for *Daphnia magna*, *Pimephales promelas* and *Algae*. Chemical classes known to be structurally similar were found to be grouped together in the TMAP chemical space, while heterogeneous classes were found to be sparse. Data imputation allowed for the identification of known and potential chemical mechanisms of action. Indeed, acethylcolinesterase and transthyretin were confirmed as major mechanisms of action of organothiophosphate and brominated flame retardant toxicity in *Daphnia magna* and *Pimephales promelas*, respectively. Overall, transdisciplinary toxicological databases combined with TMAP, stand out as a powerful, fast and scalable method to explore large datasets, allowing for meaningful and interpretable associations between chemical structures and chemical hazard identification.

**Graphical Abstract:** 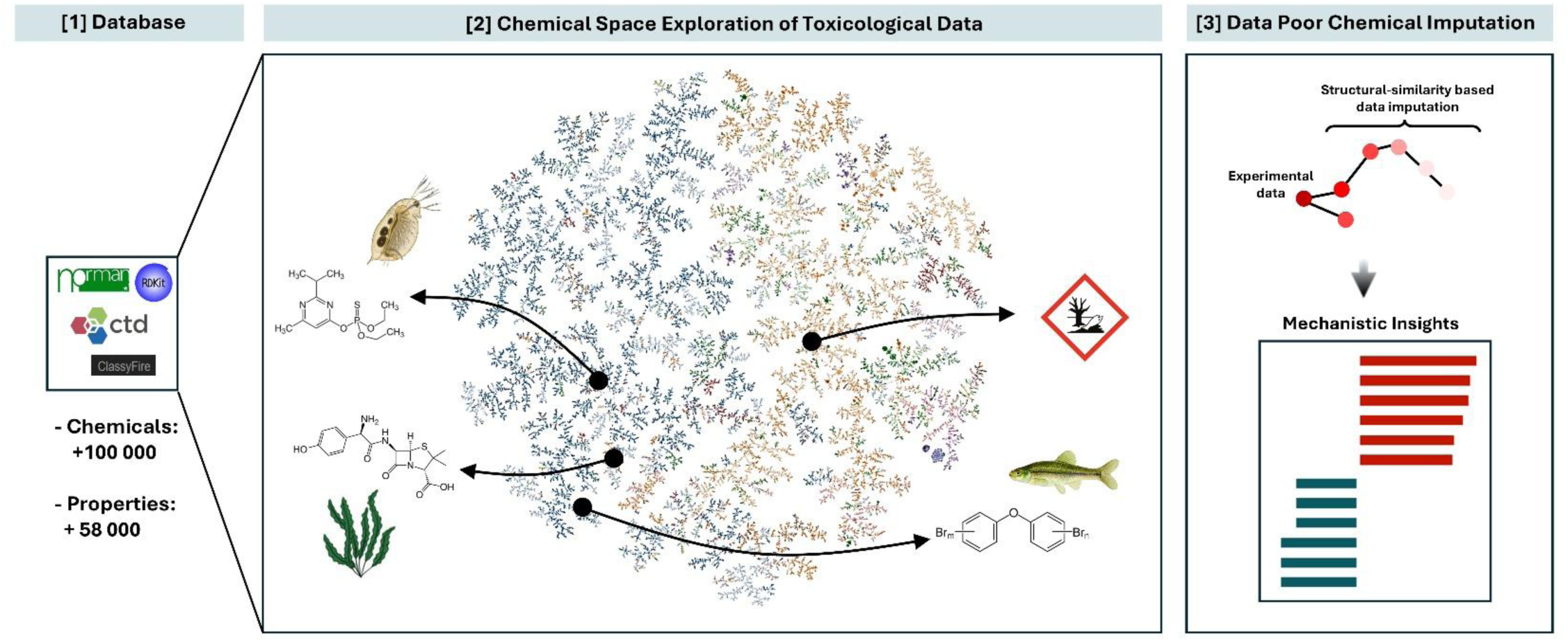

**Highlights:**

– Transdisciplinary toxicological datasets are unique resources for hazard identification.
– TMAP is a powerful tool for the visualization of very large ecotoxicological datasets.
– Graph-based imputation helps to identify targets for additional toxicological evaluation.

## INTRODUCTION

Chemical pollution affects human health and ecosystems, largely contributing to the current increase in chronic diseases and to the worldwide biodiversity loss (Malaj et al., 2014; Persson et al., 2022; Posthuma et al., 2019; Sigmund et al., 2023; van de Meent et al., 2020). For the environment, chemical pollution, interferes with ecosystem homeostasis and disrupts the food chain, leading to the irreversible shrinking of birds (Grier, 1982; Oaks et al., 2004), amphibians (Brühl et al., 2013; Egea-Serrano et al., 2012), fish populations (Brain & Prosser, 2022; Martínez-Gómez & Vethaak, 2019) and macroinvertebrates (Heß et al., 2024; Nguyen et al., 2023) across the globe.

With over 350 000 chemicals and mixtures currently registered for production and use worldwide (Wang et al., 2020) and new compounds being introduced at an exponential rate (Llanos et al., 2019), the need to identify and characterize chemical hazards has never been more urgent. The European Environment Agency (EEA) estimates that up to 90% of authorized chemicals lack adequate ecotoxicity data, leaving most chemicals poorly characterized (EEA-SOER, 2020; Kristiansson et al., 2021). Despite regulatory efforts, assessing the hazards of such a large number of chemicals remains a daunting challenge. *In vivo* testing, while a gold standard for assessing ecotoxicity is slow, costly, and relies on animal models (Pastorino et al., 2024; Price et al., 2022). As a result, there is an urgent need for high-throughput, cost-efficient, and reliable screening methods, while also reducing the need for unnecessary animal experimentation. With the creation of institutional programs, rendering available large datasets of chemicals and biological properties, such as NORMAN (www.norman-network.com) (Mohammed Taha et al., 2022) and CompTox (https://comptox.epa.gov/dashboard) (Williams et al., 2017), computational methods have the potential to stand out for their capacity to generate predictions, allowing for the simultaneous exploration of large amounts of substances. Although numerous *in silico* New Alternative Methodologies (NAMs) have been developed, these models are often trained on highly stratified data, relying on a limited range of chemical properties and ecotoxicological effects (Madden et al., 2020; Patterson et al., 2021a, 2021b; Wittwehr et al., 2020). This results in restricted applicability or need to combine multiple models for effective use. To extend its applicability, *in silico* methods may require to be sustained on a large and heterogeneous number of chemicals and properties.

In this context, we generated an innovative method for the visualization and prediction of ecotoxicological data, allowing us to obtain meaningful insights into very large and complex datasets and obtain an understanding of potential chemical mechanisms of action. To do so, Tree Manifold Approximation and Projection (TMAP), an AI-based dimensionality reduction algorithm (Probst & Reymond, 2020), were implemented combined with a large database of ecotoxicological and toxicogenomic data. Furthermore, a graph-based data imputation approach was applied as a read-across tool to identify candidate chemical targets for three different taxa relevant to ecotoxicity.

## MATERIALS AND METHODS

### Data collection and curation

A large toxicological database was built using multiple resources (Figure 1A-B). This process was achieved in various steps. (**A**) *Selection of environmentally relevant chemicals*: the NORMAN-SLE-S0.0.5.1 database (Mohammed Taha et al., 2022), commonly called Norman SusDat, containing more than 120 000 structures was selected. This database integrates chemicals from a myriad of chemical classes (eg. Pharmaceuticals, parabens, PFAS) and contains valuable information such as cross-platform identifiers (e.g. InChIKey, CAS Registry numbers, European Community EC numbers, PubChem ID) and SMILES. (**B**) *Ecotoxicological data*: NORMAN SusDat contains acute experimental and predicted toxicity values for four different taxa: the microcrustacea *Daphnia magna*, the ciliated protozoan *Tetrahymena pyriformis*, *Algae (Selenastrum capricornutum)* and the fish *Pimephales promelas*. Predicted acute toxicity values were generated by the QSAR model described in (Aalizadeh et al., 2017). (**C**) *Toxicogenomics data*: chemical-gene/protein and chemical-pathway/phenotype interactions were retrieved from the Comparative Toxicogenomics Database (https://ctdbase.org) (Davis et al., 2025). Only chemicals present in NORMAN SusDat were selected. (**D**) *Physicochemical information*. Physicochemical properties, including molecular mass, logKow, vapor pressure and polarity were computed for each chemical using *rdkit* (v2024.3.5) in Python (v3.14) based on SMILES. (**F**) *Fingerprints*. Using SMILES, MHFP6 fingerprints (Probst & Reymond, 2018) were computed in Python, allowing chemical substructures to encode in a 2048 bits vector. (**G)** *Chemical Taxonomy*. Classyfire (Djoumbou Feunang et al., 2016) was used to generate a chemical taxonomy for all chemicals present in the database based on *InchKey* identifiers. Classyfire generates an automated hierarchical chemical classification, providing for each compound a broad chemical classification (e.g. organic, inorganic compounds) and a general chemical class (e.g. lipid and lipid-like molecules) and multiple subclasses (e.g. fatty acyls) attribution. Using NORMAN SusDat, 97.2% of the substances were successfully classified, remaining were set as unknowns. The database was assembled and curated using R (v4.5.0) and Python.

**Figure 1.**
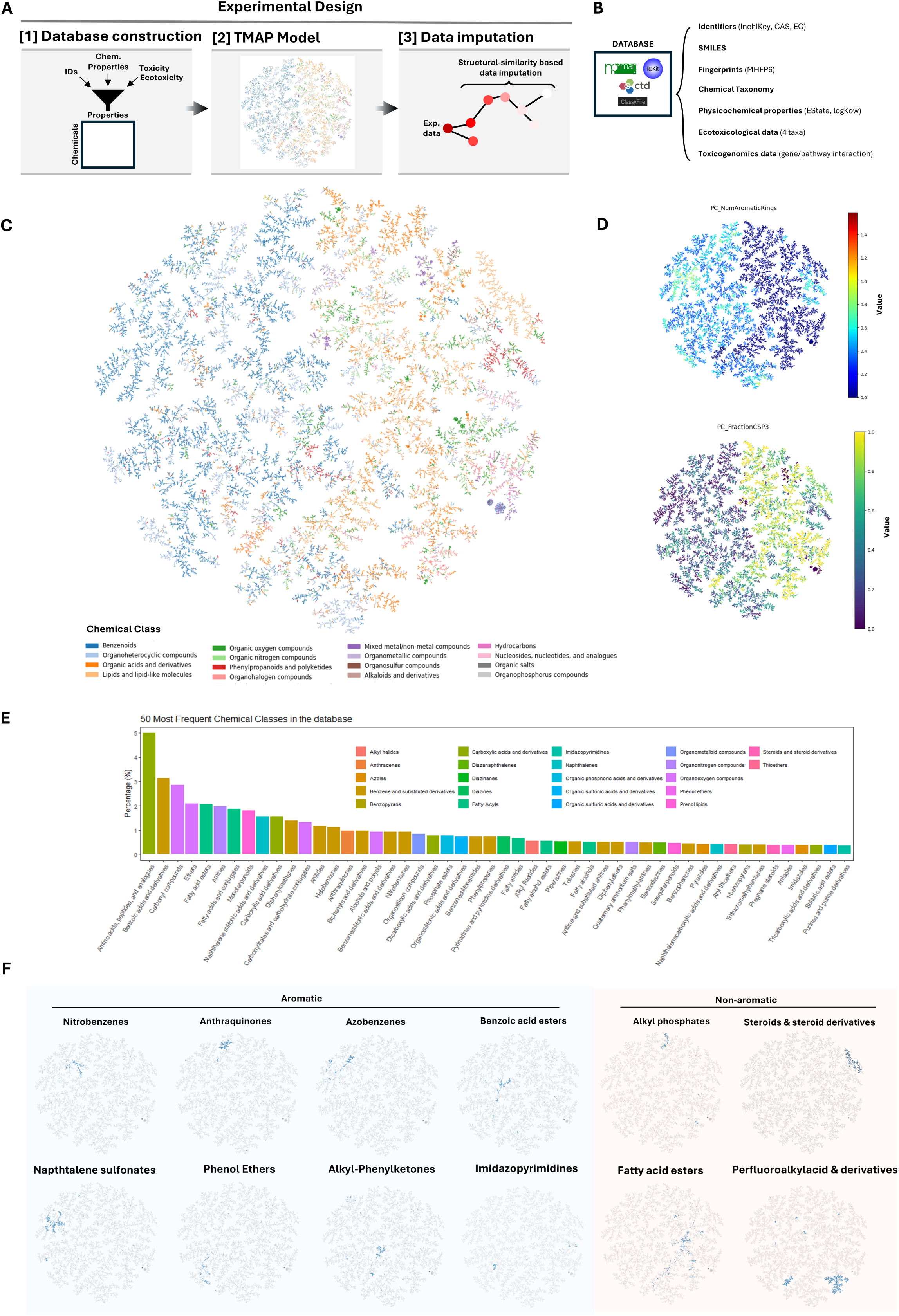
Experimental design and description of the chemical space. (**A**) A transdisciplinary chemical database was used to generate a TMAP model of the chemical space. TMAP embeddings were used to develop a data imputation model, used for the exploration of ecotoxicological data. (**B**) The database was built by combining information from multiple resources, including NORMAN (identifiers, SMILES, predicted ecotoxicological data), RDKIT (physicochemical properties and fingerprints), CTD (toxogenomics data) and Classyfire (chemical taxonomy). (**C**) TMAP chemical space representing the top chemical classes in the dataset. Nodes are chemicals and edges represent chemical structural similarities. (**D**) TMAP representation of the number of aromatic rings (top) and the fraction of aliphatic (sp³) carbons. (**E**) Barplot representing the 50 most frequent chemical classes in the database using Classyfire taxonomy. (**F**) Representative illustration of aromatic and non-aromatic chemical subclasses in the chemical space.

### TMAP dimensionality reduction model

To organize chemicals according to their structural similarity, a dimensionality reduction model was implemented. TMAP (v 1.0.6) is based on LSH forest and *k*-nearest neighbor (*k-NN*) approximation combined with minimum spanning tree (MST) computation based on the Kruskal’s algorithm, allowing to obtain a fast visualization of large datasets of chemical based on structural similarity (Probst & Reymond, 2020). TMAP was implemented in Python using MHFP6 fingerprints and was optimized using the following parameters: d=512, l=128, k=15, kc=10, p=1/50. Further details and an explanation of the rationale and the impact of each parameter on the model can be found in Probst & Reymond, 2020.

### Data imputation

A graph-based diffusion method based on TMAP chemical space was developed to impute property values for data-poor chemicals in the database. The goal of this imputation model is to estimate data gaps for chemicals based on their graph connectivity. Indeed, the TMAP graph *G=(V,E)* represents chemicals as nodes (*V*) and relationships (structural similarity) as edges (*E*). For each chemical property in the database, if a chemical has a known value for a given feature, the algorithm propagates this value to its neighboring nodes within a given depth, weighted by their graph distance. The depth limit (*d*) is used for breadth-first search. For each chemical *i* with an experimental value *v*i, each neighbor *j* at a distance *dist(i,j)* will obtain a propagation value as 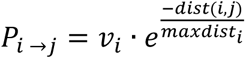 where *maxdist* is the maximum distance within the subgraph reached by the breadth-first search from *i*, limited by the depth limit *d*. Thus, the propagation uses an exponential decay function where closer neighbors receive higher weights than distant ones. After propagating values from all nodes that have observed data, the algorithm aggregate propagated contributions as 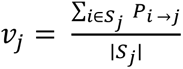 where *v_j_* represent imputed values from the propagation and *S_j_* is the set of source nodes that propagated a value to *j*. Depth (*d*) was conservatively fixed to 5. If a node already had an observed value, it remains unchanged in the final output matrix.

## RESULTS AND DISCUSSION

### A dimensionality reduction model of environmental-relevant chemicals

A comprehensive database was generated through the integration and curation of data from multiple transdisciplinary sources (Figure 1A-B). From the original 120 530 chemicals in the NORMAN SusDat database, 10 887 were removed following curation and fingerprint computation. The final database contains 109 643 chemicals and 58 544 chemical properties, including physicochemical properties (n = 213), toxicogenomics data (n = 58 327) and predicted acute ecotoxicological data (n = 4) (Figure 1B).

Given the high dimensionality of the database, TMAP was used as a dimensionality reduction model to obtain latent variables representing chemical structural similarities. Indeed, in TMAP embeddings, chemicals proximity and vertex connectivity represent the structural similarity between the compounds. Similar groups of compounds are expected to be represented within the same sub-branches, closely associated. This model has been successfully implemented in the field of drug discovery and the classification of natural products (Moine-Franel et al., 2024; Rutz et al., 2022). The chemical space (Figure 1C) shows a clear separation between major chemical classes, that can be explained by their physicochemical properties. In blue, on the left side, we can find a predominance of benzenoid compounds. On the other side, in orange, we can find predominantly lipids and lipid-like molecules. At a broader scale, TMAP grouped chemicals based on the presence or absence of aromatic structures and according to their chemical complexity, which can be clearly seen in Figure 1D. Benzenoids and most organoheterocyclic compounds are predominantly aromatic, featuring electron-rich *π*-systems that contribute to moderate-high polarity, especially when heteroatoms are present (Figure 1C–D). In contrast, organic acids, lipid and lipid-like molecules tend to have high aliphatic (*sp³*) carbon content, indicating greater complexity and hydrophobicity (Figure 1D).

The most frequent chemical class in the database were carboxylic acids and derivatives, including amino acids, peptides and analogs (Figure 1E). This was followed by benzenes (e.g. diphenylmethanes and anilides) and organooxygen compounds (e.g. carbonyl compounds, ethers). At a lower-level chemical taxonomy, TMAP clustered compounds within subclasses characterized by a known strong chemical similarity. As a proof of concept, group of chemicals were highlighted in Figure 1F. Aromatic chemicals such as nitrobenzenes, anthraquinones, azobenzenes, benzoic acids, naphthalenes, phenol ethers, and imidazopyrimidines were highly clustered, as were non-aromatic compounds like alkyl phosphates, fatty acid esters, perfluorinated compounds and steroids (Figure 1F). Steroid and steroid derivatives (e.g. estradiol benzoate, aldosterone, pregnenolone) are characterized by a high number of aliphatic carbocycles, property clearly defined in the chemical space (Figure 1F, Figure S1A). In addition, the strong perfluorinated compound clustering can be explained by multiple physicochemical properties. Indeed, they display very low minimum electrotopological state index values due to highly electron-deficient atoms from multiple fluorines and have moderate monoisotopic mass, are enriched in heavy and heteroatoms and show moderate to high molecular complexity (i.e low Bertz and Balaban J indexes) (Figure S1B-G).

In contrast, more heterogeneous chemical subclasses such as alkyl-phenylketones and fatty acid esters are widely distributed in the chemical space (Figure 1F). This broad dispersion can be explained by their structural diversity, which includes variations in alkyl chain length, functional groups and aromatic substitutions. These variations lead to greater variability in physicochemical properties such as lipophilicity, molecular weight, polarity, and topological indices, resulting in less cohesive clustering compared to the highly uniform structures of perfluorinated compounds and others.

In summary, the TMAP chemical space displays an accurate and meaningful visual representation of chemicals distributed according to their structural similarity, allowing for the exploration of chemical properties and toxicological data. Heterogeneous chemical categories are sparse, while known strongly homogeneous classes are consistently clustered.

### Chemical space of bioaccumulative chemicals

To explore the relevance of our database and TMAP chemical space for chemical hazard identification, we evaluated the bioaccumulation distribution within the chemical space. Chemicals with high bioaccumulation potential here are defined as having a logKow > 3. According to the EU Registration, Evaluation, Authorisation and Restriction of Chemicals (REACH) Annex IX for substances registered at ≥100 tons/year, additional testing for long-term aquatic toxicity and bioaccumulation potential may be required depending on, among other factors, logKow. Indeed, substances with a logKow > 3 are considered to have a potential for bioaccumulation, which can trigger further assessment, including the possibility of a fish bioaccumulation study if no alternative data (e.g., QSAR, *in vitro*) are available (REACH Regulation, 2025). Figure 2A shows that high bioaccumulative compounds, represented in dark brown, are grouped in specific regions of the TMAP. Three major groups can be highlighted. In the top left side, we can observe a highly grouped cluster of complex highly bioaccumulative chemicals, mostly substituted aniline derivatives, with amide, carbamate, azo and pyrazolone moieties (e.g. 2-(3-Pentadecylphenoxy)ethyl[4-[[4,5-dihydro-4-[(4-methoxyphenyl)azo]-5-oxo-1-(2,4,6-trichlorophenyl)-1H-pyrazol-3-yl]amino]phenyl]carbamate). These chemical features are relevant from a regulatory and toxicological perspective because they are frequently associated with persistent, bioaccumulative and toxic (PBT) profiles. Azo and aniline derivatives are widely associated with carcinogenic or mutagenic effects in aquatic organisms and mammals (DeMarini et al., 2020; Feng et al., 2012). In the middle square, aryl organophosphate triesters (e.g. TBEP, TNPP), compounds used as organophosphate flame retardants can be found. These compounds are known for their persistence and bioaccumulation potential, neurotoxic effects, and endocrine-disrupting properties (van der Veen & de Boer, 2012; J. Yang, Zhao, et al., 2019). Finally, on the right side, we observe a large and diverse category of bioaccumulative fatty acyl compounds, including lubricants such as (di)pentaerythritol esters (e.g. Pentaerythritol tetrabehenate, Dipentaerythritol pentastearate), polyol-based emollients commonly used in cosmetics and skin care (e.g. Tridocosyl citrate), methacrylates and acrylates (e.g., Hexadecyl icosyl methacrylate, Hexadecyl icosyl acrylate) and maleate derivatives used as plasticizers and crosslinkers (e.g., Bis[[tetrahydro-4-hydroxy-3,5,5-tris[[(1-oxooctadecyl)oxy]methyl]-2H-pyran-3-yl]methyl] maleate), among many others.

**Figure 2.**
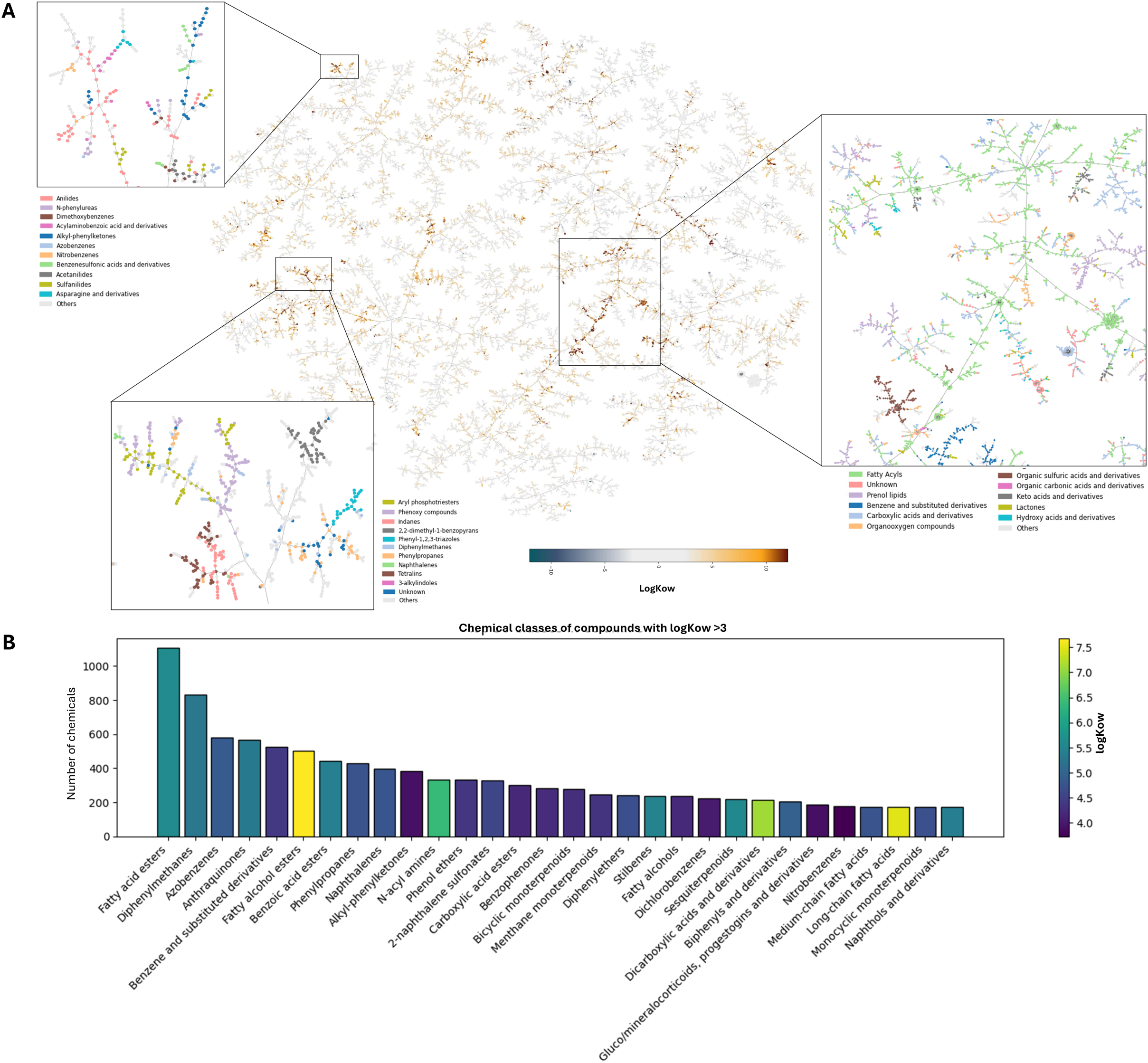
Exploration of chemical bioaccumulative potential. (**A**) Representation of *logKow* values in the chemical space and zoom at three particular regions, shown the taxonomical classification of the compounds. (**B**) Barplot representing the most frequent chemical classes having a *logKow* > 3. Colors represent the average logKow within the chemical class.

Indeed, fatty acyls and their derivatives appear to be the most predominant bioaccumulative category in our database (Figure 2B). Fatty alcohol esters are the category of chemicals displaying the highest average logKow. Aromatic compounds as substituted benzenes and azoles were also highly represented among chemicals with high logKow.

As an exploratory tool, TMAP clusters and reliably represent known chemical classes associated with bioaccumulation, providing a visual framework for identifying and prioritizing substances of concern and eventually guiding subsequent toxicological endpoint assessments.

### Chemical Space of *Daphnia magna* Toxicity

To evaluate the relevance of TMAP for aquatic ecotoxicology, we next evaluated the distribution of chemicals toxic to microcrustacea *Daphnia magna*. Figure 3A-B highlights the groups of compounds identified as the most toxic. The most abundant chemical classes associated with *Daphnia magna* toxicity are organofluorides and organoiodides (Figure 3A). Their close association in the chemical space reflect their chemical structural similarity and shared physicochemical profiles, such as high molecular mass and halogenation, factors associated with bioaccumulation and toxicity (Figure 3B, Figure S2A-B). Organofluorine compounds are widely used as components for materials, agrochemicals and pharmaceuticals (Logeshwaran et al., 2021a; Prakash, 2015). Per- and polyfluoroalkyl substances, known as PFAS, are recognized to be chemical of concern for the aquatic life (Ahrens & Bundschuh, 2014). In particular, PFOS has been shown to induce acute and long-term toxicity on *Daphnia at* environmental concentrations (Jeong et al., 2016; Logeshwaran et al., 2021b; H.-B. Yang, Zhao, et al., 2019). Organoiodides are used as disinfectants and pesticides. In particular, perfluorinated alkyl iodides, compounds with a fully fluorinated carbon chains of varying lengths with an iodine substituent at a mid-chain position, representing a prominent class of compounds strongly associated with predicted *Daphnia magna* toxicity. Representative examples are henicosafluoro-, pentacosafluoro-, and other perfluoroalkyl chains with iodine atoms typically positioned between carbons 9 to 14 (e.g. Henicosafluoro-12-iodononadecane, Pentacosafluoro-14-iodoicosane) (Figure S2C).

**Figure 3.**
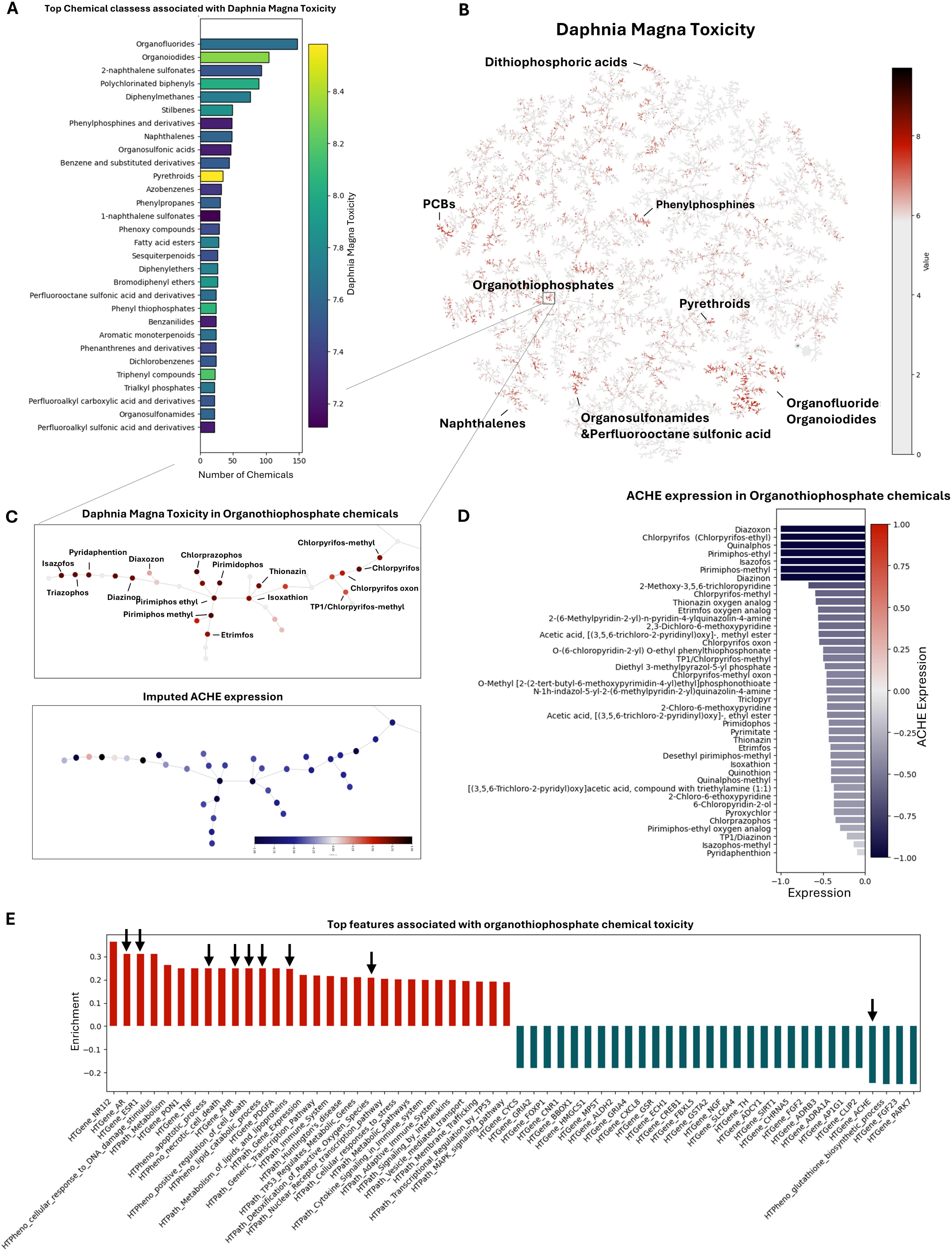
Exploration of *Daphnia magna* toxicity. (**A**) Representation of the most frequent chemical classes associated with *Daphnia magna* toxicity. Colors represent the average toxicity values per class. (**B**) TMAP representation of *Daphnia magna* toxicity with illustrative examples. Zoom on the organothiophosphate chemical branch, displaying *Daphnia magna* toxicity (top) and imputed ACHE expression (bottom). (**D**) Chemicals displaying the lowest ACHE expression in the organothiophosphate chemical branch. (**E**) Top upregulated (red) and downregulated (blue) features associated with organothiophosphate chemical toxicity. ACHE: acethylcolinesterase.

From the chemical taxonomy, the most toxic class for *Daphnia magna* appears to be the pyrethroids (Figure 3A). Pyrethroids are used as insecticides in agriculture, but also for pets or against mosquitoes. These synthetic and strongly structurally associated insecticides form a cluster within the chemical space (Figure 3B). Indeed, these compounds have a cyclopropane ring fused to a phenoxybenzyl (Matsuo, 2019) and are known to display acute toxicity in *Daphnia* (Mokry & Hoagland, 1990). Among them, Acrinathrin, Tralomethrin and the well-known Permethrin were found (Figure S2D).

Another family toxic to *Daphnia magna*, organothiophosphates, are also highly clustered (Figure 3A). This family includes compounds such as chlorpyrifos, an insecticide used in agriculture for decades and known to be toxic for aquatic species, especially macroinvertebrates (X. Huang et al., 2020). To test the potential of TMAP to explore and infer chemical mechanisms of action, toxicogenomics data was combined with a graph-based data imputation method. For organothiophosphates, acetylcholinesterase (ACHE) inhibition is a major well described mechanism of action of these compounds, together with noncholinergic mechanisms of neurotoxicity (Tsai & Lein, 2021). Exploring the data in the chemical space combined with the imputation allowed us to observe that chlorpyrifos, together with most chemicals in the organothiophosphate family has strong ACHE inhibition, as expected (Figure 3C) (Tsai & Lein, 2021). Beyond this known mechanism of action, the imputation approach allows identifying additional genes up- or downregulated based on human and animal model experimental data (Figure 3E). For example, the increase in signals associated with lipid metabolism, oxidative stress and cell death and the strong upregulation of androgen receptor (AR), estrogen receptor 1 (ESR1) and aryl hydrocarbon receptor (AHR). This might suggest disrupted lipid homeostasis, compromised cellular stability and potential endocrine-disrupting effects of organothiophosphates, which has been suggested by others (Zhang et al., 2024). Even if not directly transferable to *Daphnia magna*, microcrustacea having differences in endocrine system compared to mammals (Cho et al., 2022), such a comparison highlights potential endocrine compounds for the aquatic life (Yang, Li, et al., 2019).

### Chemical Space of *Pimephales Promelas* Toxicity

The fish *Pimephales Promelas* appears to be more sensitive to anthraquinone dyes (e.g. 1,5-Diaminoanthraquinone, Solvent Blue 58) (Figure 4A-B, Figure S3A-C) and benzoic acid esters (including phthalates as Bis(2-ethylhexyl) tetrabromophthalate, TBPH) (Figure S3D-E), followed by organofluorides, similarly to *Daphnia magna*. Anthraquinone dyes are widely used in the textile and cosmetic industries and are characterized by high chemical stability, high hydrophobicity and low biodegradability, suggesting a high toxicity for aquatic environments (Farias et al., 2023). Exposure to anthraquinone dyes has been associated with reproductive toxicity in *Pimephales Promelas (Parrott et al., 2016)*. In turn, benzoic acid esters as the flame retardant TBPH are structurally complex and resistant to biodegradation, representing a concern for environmental persistence and trophic transfer. TBPH has been shown to induce genotoxicity in *Pimephales Promelas (Bearr et al., 2010)*.

**Figure 4.**
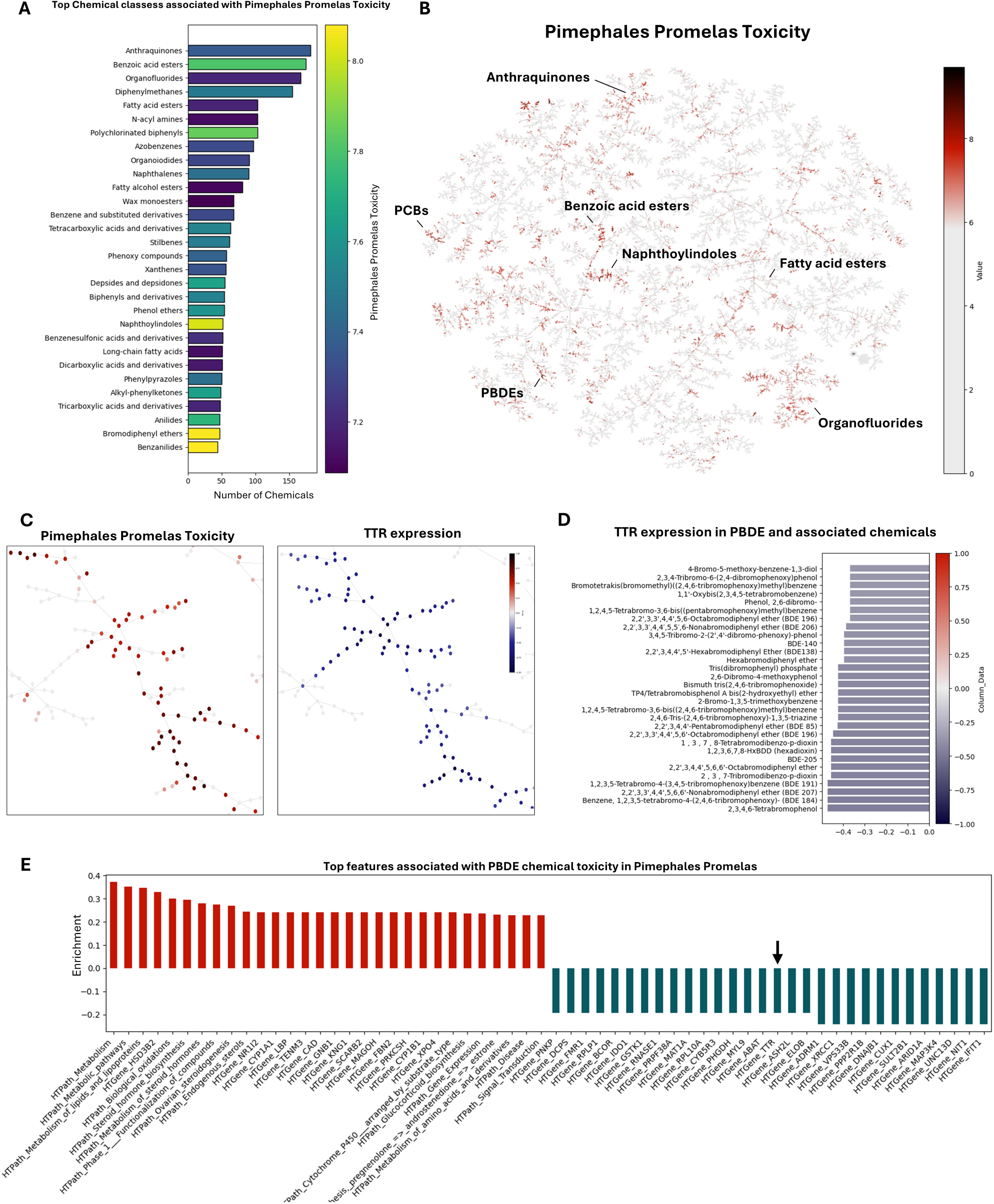
Exploration of *Pimephales Promelas* toxicity. (**A**) Representation of the most frequent chemical classes associated with *Pimephales Promelas* toxicity. Colors represent the average toxicity values per class. (**B**) TMAP representation of *Pimephales Promelas* toxicity with illustrative examples. (C) Zoom on the PBDE chemical branch, displaying *Pimephales Promelas* toxicity (left) and imputed TTR expression (right). (**D**) Chemicals displaying the lowest TTR expression in the PBDE chemical branch. (**E**) Top upregulated (red) and downregulated (blue) features associated with PBDE chemical toxicity. TTR: Transthyretin. PBDE: Polybrominated diphenyl ethers.

**Figure 5.**
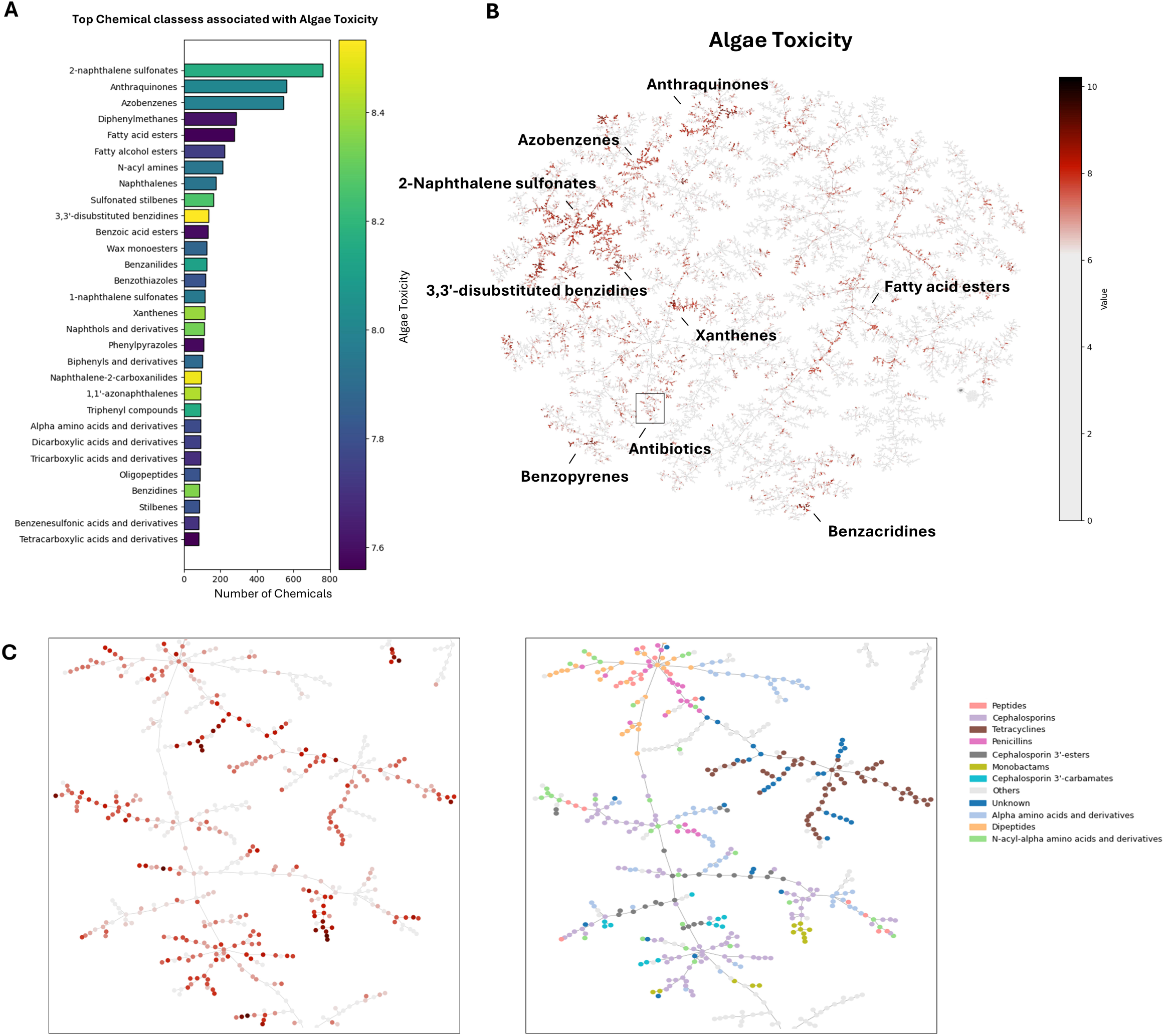
Exploration of *Algae* toxicity. (**A**) Representation of the most frequent chemical classes associated with *Algae* toxicity. Colors represent the average toxicity values per class. (**B**) TMAP representation of *Algae* toxicity with illustrative examples. (C) Zoom on a chemical branch containing multiple antibiotic classes, displaying *Algae* toxicity (left) and imputed chemical taxonomy (right).

Polychlorinated biphenyls (PCBs) and polybrominated diphenyl ethers (PBDEs) are two major categories of compounds strongly clustered in the TMAP that are known to be toxic to *Pimephales Promelas* are (Figure 4A-B). Multiple PCBs and congeners were found to be highly toxic in these clusters, such as Decachlorobiphenyl (PCB 209), 2,2′,3,3′,4,4′,5,5′,6-Nonachlorobiphenyl (PCB 206) and 2,2′,3,3′,4,4′,6,6′-Octachlorobiphenyl (PCB 197). In turn, PBDE flame retardants, such as 1,1’-Oxybis[2,3,4,5,6-pentabromobenzene] (BDE209), 2,2’,3,4,4’,5’,6-Heptabromodiphenyl ether (BDE 183) and 2,2’,3,3’,4,4’,5,6,6’-Nonabromodiphenyl ether (BDE 207) are highly brominated, bioaccumulative and toxic compounds. Toxicogenomics data shows that most PBDEs selectively may inhibit transthyretin (TTR) receptor expression, a thyroid hormone transport protein (Figure 4C-D), a shown previously (Meerts et al., 2000). The strongest TTR inhibitors were 2,3,4,6-Tetrabromophenol, BDE 184 and BDE 207. TTR is a carrier of thyroid hormones, critical for neurodevelopment, metabolism, and growth (Bauer et al., 2002). Thyroid glands are the primary target of PBDEs, primarily because of their strong structural similarity to endogenous thyroid hormones. Early life *Pimephales promelas* exposure to PBDEs has been associated with thyroid, hepatic and immunological disorders (Noyes et al., 2011; Thornton et al., 2018). As suggested by toxicogenomic imputed data, PBDEs affect multiple biological pathways beyond potential thyroid disruption. Disruption of CYP1A1 and CYP1B1 indicate activation of AhR-mediated xenobiotic metabolism, whilst HSD3B2 and SULT2B1 suggest steroid hormone synthesis and metabolism disruption. PBDEs AhR-mediated toxicity has been proven using zebrafish (Liu, Tang, et al., 2015). Targets of neurodevelopment and immunological alterations are TENM3, UNC13D, LBP and IFIT1. Finally, alterations on NR1L1, MAP3K4, and GNB1 expression suggest a potential impact on lipid metabolism and cell signaling. These features may serve as potential candidates to be evaluated in future studies evaluating PBDE *Pimephales promelas* toxicity.

### Algae Toxicity

*Algae* data suggest that 2-naphthalene sulfonates, anthraquinones and azobenzenes are the three major chemical classes that are the most frequently toxic. 2-naphthalene sulfonates and anthraquinones are commonly used in the industrial sector for the production of dyes and colorants (Mehra & Chadha, 2020). 2-naphthalene sulfonates have multiple sulfonate groups and azo linkages, high polarity and resistance to biodegradation. Nitro-substituted azo dyes (e.g. 2,7-Naphthalenedisulfonic acid, 4-[[4-[[[[5-hydroxy-6-[[4-[(4-nitro-2-sulfophenyl)amino]phenyl]azo]-7-sulfo-2-naphthalenyl]amino]carbonyl]amino]-5-methoxy-2-methylphenyl]azo]-5-[(phenylsulfonyl)oxy]-, sodium salt) and chlorotriazine derivatives (e.g. *Octasodium 2,2’-[(2,2’-disulphonato[1,1’-biphenyl]-4,4’-diyl)bis[imino(6-chloro-1,3,5-triazine-4,2-diyl)imino(1-hydroxy-3-sulphonatonaphthalene-6,2-diyl)azo]]bisnaphthalene-1,5-disulphonate)* are compounds particularly hazardous to aquatic life, including *Algae* (Hashemi & Kaykhaii, 2022). In turn, anthraquinones are persistent aromatic structures (high logKow), which contribute to their bioaccumulation in aquatic environments. Certain derivatives, such as 1-amino-2-bromo-4-hydroxyanthraquinone or 1,4-diaminoanthraquinone are associated with growth inhibition and photosynthetic disruption in *Algae* (Tukaj & Aksmann, 2007). Finally, azobenzenes are compounds characterized by their -N=N-azo linkage and are known photoresponsive molecules (Londoño-Berrío et al., 2022).

Further exploration of the TMAP chemical space allowed us to define antibiotics as one relevant chemical category for *Algae* toxicity. The excessive use of antibiotics makes them to be commonly detected in water systems, entering via domestic or medical facility wastewater or livestock-farming (Le et al., 2023). It has been estimated that nearly one-fifth of antibiotics may cause hazard to marine algae (Zhou et al., 2022). TMAP displayed multiple antibiotic classes situated closely in chemical space, including penicillin, cephalosporins, tetracyclines and monobactams. These compounds are structurally related through shared core scaffolds (e.g. β-lactam rings) and/or similarities in molecular descriptors (e.g. polarity, hydrogen-bonding capacity, ring topology). Among penicillin, Amoxicillin, Benzylpenicillin and Phenoxymethylpenicillin-Benzathine were found in our list of compounds toxic for Algae. Amoxicillin has been shown to impact algae growth (Liu, Chen, et al., 2015). Cephalosporins (e.g. Ceftaroline Fosamil, Cefditoren Pivoxil, Cefditoren), commonly used to treat bacterial infections (e.g. pneumonia, skin infections, urinary tract infections) are known to inhibit algal growth and promote antibiotic resistance genes (Guo et al., 2015; F.-L. Huang et al., 2024). In turn, tetracyclines (e.g. Tigecycline, Minocycline hydrochloride, Lymecycline) strongly impacts algal photosynthesis (Chen et al., 2022). Finally, monobactams (e.g. Aztreonam, Carumonam and Tigemonam) are less known toxicants for *Algae*.

These antibiotic categories are known to inhibit bacterial cell wall synthesis by preventing peptidoglycan cross-linking. In algae, Chen et al. (2022) identified photosynthetic electron transport and enzymes to be impacted by tetracycline exposure. Given the limited available toxicogenomics data for these compounds, no particular mechanism was found, suggesting the dependence on experimental studies to obtain reliable candidates.

### Strengths and Limitations of the model and Perspectives

Here we generated a transdisciplinary database and a dimensionality reduction model to explore physicochemical properties and ecotoxicity data for three different taxa relevant for aquatic toxicity, considering the chemical structure association and their potential mechanisms of action. As proof of concept, most common and frequent chemical class toxicity were discussed, exploring thus only a fraction of the capabilities of the model. The use of visualization tools that accurately represent chemicals according to their structures streamline hazard identification and serve as a tool to generate hypothesis that may be further validated with complementary methods. Indeed, the database and the TMAP model can be used to eventually find safer chemical alternatives, to find candidate chemicals of concern for further investigation, to perform read-across and to serve as a basis for the prediction of different toxicity endpoints. Available information in the database can be further complemented by adding additional repositories (e.g. docking, binding assay data, *in vitro* data). Given the extensive capabilities of the graph-based imputation model, machine learning models can be implemented for the prediction of endpoints as skin sensitization, endocrine disruption or neurotoxicity.

A major advantage of TMAP is its potential for scalability. The model was developed to have the capability to be implemented using millions of compounds, allowing for the toxicological assessment of more diverse and relevant structures (Probst & Reymond, 2020). Visualizing extremely large datasets improve interpretability, allowing to focus on the generation of hypothesis.

The approach described also comes with multiple important limitations. Graph-based imputation, although based on structural similarities, ignores mechanistic biology. The method assumes that connected chemicals share similar feature values, although in toxicology, structurally similar chemicals may sometimes have very different biological effects. To account for this, a very conservative exponential decay function and depth was applied, to avoid propagating values to structurally distinct chemicals. Future efforts may need to implement multiple decay functions corrected by rules for biological plausibility, including uncertainty imputation quantifications. Additionally, the model capabilities is limited to available toxicogenomic data. The CTD database is indeed a unique resource that is frequently updated that may required to be complemented with data from additional tools (e.g. *CompTox*). CTD lacks quantitative information, limiting the capabilities of the model as it does not consider chemical thresholds and allow to define chemical targets at low relevant concentrations. Furthermore, the divergence between database in data organization, the use of diverse identifiers and file formats makes chemoinformatic and toxicological data to be fragmented. This lead to obtention of noisy data that requires strong efforts for curation before being useful. Thus, extra curation to sort and use only reliable data is a must-do.

In conclusion, the described approach, allowing the visualization and exploration of very large toxicological datasets may have the potential to serve as a tool for the prioritization of chemicals for hazard identification and the prediction of toxicological endpoints.

## AUTHOR CONTRIBUTIONS

DLR and NC conceived the study. DLR and GGL curated the data and DLR performed the formal analysis, developed the code for data analysis and created all visualizations. Project administration, funding acquisition and drafting of the original manuscript was done by DLR and NC and all authors reviewed and edited the final version.

## ACKNOWLEDGEMENTS

The authors thank Dr. Oliver Warwick for his support and valuable collaboration in this study.

## References

Aalizadeh, R., von der Ohe, P. C., & Thomaidis, N. S. (2017). Prediction of acute toxicity of emerging contaminants on the water flea Daphnia magna by Ant Colony Optimization–Support Vector Machine QSTR models. Environmental Science: Processes & Impacts, 19(3), 438–448. 10.1039/C6EM00679E

Ahrens, L., & Bundschuh, M. (2014). Fate and effects of poly- and perfluoroalkyl substances in the aquatic environment: A review. Environmental Toxicology and Chemistry, 33(9), 1921–1929. 10.1002/etc.2663

Bauer, M., Heinz, A., & Whybrow, P. C. (2002). Thyroid hormones, serotonin and mood: of synergy and significance in the adult brain. Molecular Psychiatry, 7(2), 140–156. 10.1038/sj.mp.4000963

Bearr, J. S., Stapleton, H. M., & Mitchelmore, C. L. (2010). Accumulation and DNA damage in fathead minnows (Pimephales promelas) exposed to 2 brominated flame-retardant mixtures, Firemaster® 550 and Firemaster® BZ-54. Environmental Toxicology and Chemistry, 29(3), 722–729. 10.1002/etc.94

Brain, R. A., & Prosser, R. S. (2022). Human induced fish declines in North America, how do agricultural pesticides compare to other drivers? Environmental Science and Pollution Research, 29(44), 66010–66040. 10.1007/s11356-022-22102-z

Brühl, C. A., Schmidt, T., Pieper, S., & Alscher, A. (2013). Terrestrial pesticide exposure of amphibians: An underestimated cause of global decline? Scientific Reports, 3(1), 1135. 10.1038/srep01135

Chen, Z., Gu, G., Wang, Z., Ou, D., Liang, X., Hu, C., & Li, X. (2022). Effects of tetracycline on Scenedesmus obliquus microalgae photosynthetic processes. International Journal of Molecular Sciences, 23(18), 10544.

Cho, H., Ryu, C. S., Lee, S.-A., Adeli, Z., Meupea, B. T., Kim, Y., & Kim, Y. J. (2022). Endocrine-disrupting potential and toxicological effect of para-phenylphenol on Daphnia magna. Ecotoxicology and Environmental Safety, 243, 113965. 10.1016/j.ecoenv.2022.113965

Davis, A. P., Wiegers, T. C., Sciaky, D., Barkalow, F., Strong, M., Wyatt, B., Wiegers, J., McMorran, R., Abrar, S., & Mattingly, C. J. (2025). Comparative Toxicogenomics Database’s 20th anniversary: update 2025. Nucleic Acids Research, 53(D1), D1328–D1334. 10.1093/nar/gkae883

DeMarini, D. M., Carreón-Valencia, T., Gwinn, W. M., Hopf, N. B., Sandy, M. S., Bahadori, T., Calaf, G. M., Chen, G., de Conti, A., Fritschi, L., Gi, M., Josephy, P. D., Kirkeleit, J., Kjaerheim, K., Langouët, S., McElvenny, D. M., Sergi, C. M., Stayner, L. T., Toyoda, T., … Schubauer-Berigan, M. K. (2020). Carcinogenicity of some aromatic amines and related compounds. The Lancet Oncology, 21(8), 1017–1018. 10.1016/S1470-2045(20)30375-2

Djoumbou Feunang, Y., Eisner, R., Knox, C., Chepelev, L., Hastings, J., Owen, G., Fahy, E., Steinbeck, C., Subramanian, S., Bolton, E., Greiner, R., & Wishart, D. S. (2016). ClassyFire: automated chemical classification with a comprehensive, computable taxonomy. Journal of Cheminformatics, 8(1), 61. 10.1186/s13321-016-0174-y

EEA-SOER. (2020). The European environment-state and outlook 2020 Knowledge for transition to a sustainable Europe. 10.2800/96749

Egea-Serrano, A., Relyea, R. A., Tejedo, M., & Torralva, M. (2012). Understanding of the impact of chemicals on amphibians: a meta-analytic review. Ecology and Evolution, 2(7), 1382–1397. 10.1002/ece3.249

Farias, N. O. de, Albuquerque, A. F. de, dos Santos, A., Almeida, G. C. F., Freeman, H. S., Räisänen, R., & Umbuzeiro, G. de A. (2023). Is natural better? An ecotoxicity study of anthraquinone dyes. Chemosphere, 343, 140174. 10.1016/j.chemosphere.2023.140174

Feng, J., Cerniglia, C. E., & Chen, H. (2012). Toxicological significance of azo dye metabolism by human intestinal microbiota. FBE, 4(2), 568–586.

Grier, J. W. (1982). Ban of DDT and Subsequent Recovery of Reproduction in Bald Eagles. Science, 218(4578), 1232–1235. 10.1126/science.7146905

Guo, R., Xie, W., & Chen, J. (2015). Assessing the combined effects from two kinds of cephalosporins on green alga (Chlorella pyrenoidosa) based on response surface methodology. Food and Chemical Toxicology, 78, 116–121. 10.1016/j.fct.2015.02.007

Hashemi, S. H., & Kaykhaii, M. (2022). Chapter 15 - Azo dyes: Sources, occurrence, toxicity, sampling, analysis, and their removal methods. In T. Dalu & N. T. Tavengwa (Eds.), Emerging Freshwater Pollutants (pp. 267–287). Elsevier. 10.1016/B978-0-12-822850-0.00013-2

Heß, S., Hof, D., Oetken, M., & Sundermann, A. (2024). Macroinvertebrate communities respond strongly but non-specifically to a toxicity gradient derived by effect-based methods. Environmental Pollution, 356, 124330. 10.1016/j.envpol.2024.124330

Huang, F.-L., Qin, L.-T., Mo, L.-Y., Zeng, H.-H., & Liang, Y.-P. (2024). Mechanism of the Synergistic Toxicity of Ampicillin and Cefazoline on Selenastrum capricornutum. Toxics, 12(3), 217.

Huang, X., Cui, H., & Duan, W. (2020). Ecotoxicity of chlorpyrifos to aquatic organisms: A review. Ecotoxicology and Environmental Safety, 200, 110731. 10.1016/j.ecoenv.2020.110731

Jeong, T.-Y., Yuk, M.-S., Jeon, J., & Kim, S. D. (2016). Multigenerational effect of perfluorooctane sulfonate (PFOS) on the individual fitness and population growth of Daphnia magna. Science of The Total Environment, 569–570, 1553–1560. 10.1016/j.scitotenv.2016.06.249

Kristiansson, E., Coria, J., Gunnarsson, L., & Gustavsson, M. (2021). Does the scientific knowledge reflect the chemical diversity of environmental pollution? – A twenty-year perspective. Environmental Science & Policy, 126, 90–98. 10.1016/j.envsci.2021.09.007

Le, V. Van, Tran, Q.-G., Ko, S.-R., Lee, S.-A., Oh, H.-M., Kim, H.-S., & Ahn, C.-Y. (2023). How do freshwater microalgae and cyanobacteria respond to antibiotics? Critical Reviews in Biotechnology, 43(2), 191–211.

Liu, Chen, X., Zhang, J., & Gao, B. (2015). Hormesis Effects of Amoxicillin on Growth and Cellular Biosynthesis of Microcystis aeruginosa at Different Nitrogen Levels. Microbial Ecology, 69(3), 608–617. 10.1007/s00248-014-0528-9

Liu, H., Tang, S., Zheng, X., Zhu, Y., Ma, Z., Liu, C., Hecker, M., Saunders, D. M. V, Giesy, J. P., Zhang, X., & Yu, H. (2015). Bioaccumulation, Biotransformation, and Toxicity of BDE-47, 6-OH-BDE-47, and 6-MeO-BDE-47 in Early Life-Stages of Zebrafish (Danio rerio). Environmental Science & Technology, 49(3), 1823–1833. 10.1021/es503833q

Llanos, E. J., Leal, W., Luu, D. H., Jost, J., Stadler, P. F., & Restrepo, G. (2019). Exploration of the chemical space and its three historical regimes. Proceedings of the National Academy of Sciences, 116(26), 12660–12665. 10.1073/pnas.1816039116

Logeshwaran, P., Sivaram, A. K., Surapaneni, A., Kannan, K., Naidu, R., & Megharaj, M. (2021a). Exposure to perfluorooctanesulfonate (PFOS) but not perflurorooctanoic acid (PFOA) at ppb concentration induces chronic toxicity in Daphnia carinata. Science of The Total Environment, 769, 144577. 10.1016/j.scitotenv.2020.144577

Logeshwaran, P., Sivaram, A. K., Surapaneni, A., Kannan, K., Naidu, R., & Megharaj, M. (2021b). Exposure to perfluorooctanesulfonate (PFOS) but not perflurorooctanoic acid (PFOA) at ppb concentration induces chronic toxicity in Daphnia carinata. Science of The Total Environment, 769, 144577. 10.1016/j.scitotenv.2020.144577

Londoño-Berrío, M., Pérez-Buitrago, S., Ortiz-Trujillo, I. C., Hoyos-Palacio, L. M., Orozco, L. Y., López, L., Zárate-Triviño, D. G., Capobianco, J. A., & Mena-Giraldo, P. (2022). Cytotoxicity and genotoxicity of azobenzene-based polymeric nanocarriers for phototriggered drug release and biomedical applications. Polymers, 14(15), 3119.

Madden, Judith C, Enoch, Steven J, Paini, Alicia, & Cronin, Mark T D. (2020). A Review of In Silico Tools as Alternatives to Animal Testing: Principles, Resources and Applications. Alternatives to Laboratory Animals, 48(4), 146–172. 10.1177/0261192920965977

Malaj, E., von der Ohe, P. C., Grote, M., Kühne, R., Mondy, C. P., Usseglio-Polatera, P., Brack, W., & Schäfer, R. B. (2014). Organic chemicals jeopardize the health of freshwater ecosystems on the continental scale. Proceedings of the National Academy of Sciences, 111(26), 9549–9554. 10.1073/pnas.1321082111

Martínez-Gómez, C., & Vethaak, A. D. (2019). Understanding the impact of chemicals on marine fish populations: the need for an integrative approach involving population and disease ecology. Current Opinion in Environmental Science & Health, 11, 71–77. 10.1016/j.coesh.2019.08.001

Matsuo, N. (2019). Discovery and development of pyrethroid insecticides. Proceedings of the Japan Academy, Series B, 95(7), 378–400. 10.2183/pjab.95.027

Meerts, I. A. T. M., van Zanden, J. J., Luijks, E. A. C., van Leeuwen-Bol, I., Marsh, G., Jakobsson, E., Bergman, Å., & Brouwer, A. (2000). Potent Competitive Interactions of Some Brominated Flame Retardants and Related Compounds with Human Transthyretin in Vitro. Toxicological Sciences, 56(1), 95–104. 10.1093/toxsci/56.1.95

Mehra, S., & Chadha, P. (2020). Bioaccumulation and toxicity of 2-naphthalene sulfonate: an intermediate compound used in textile industry. Toxicology Research, 9(2), 127–136. 10.1093/toxres/tfaa008

Mohammed Taha, H., Aalizadeh, R., Alygizakis, N., Antignac, J.-P., Arp, H. P. H., Bade, R., Baker, N., Belova, L., Bijlsma, L., Bolton, E. E., Brack, W., Celma, A., Chen, W.-L., Cheng, T., Chirsir, P., Čirka, Ľ., D’Agostino, L. A., Djoumbou Feunang, Y., Dulio, V., … Schymanski, E. L. (2022). The NORMAN Suspect List Exchange (NORMAN-SLE): facilitating European and worldwide collaboration on suspect screening in high resolution mass spectrometry. Environmental Sciences Europe, 34(1), 104. 10.1186/s12302-022-00680-6

Moine-Franel, A., Mareuil, F., Nilges, M., Ciambur, C. B., & Sperandio, O. (2024). A comprehensive dataset of protein-protein interactions and ligand binding pockets for advancing drug discovery. Scientific Data, 11(1), 402. 10.1038/s41597-024-03233-z

Mokry, L. E., & Hoagland, K. D. (1990). Acute toxicities of five synthetic pyrethroid insecticides to Daphnia magna and Ceriodaphnia dubia. Environmental Toxicology and Chemistry, 9(8), 1045–1051. 10.1002/etc.5620090811

Nguyen, H. H., Welti, E. A. R., Haubrock, P. J., & Haase, P. (2023). Long-term trends in stream benthic macroinvertebrate communities are driven by chemicals. Environmental Sciences Europe, 35(1), 108. 10.1186/s12302-023-00820-6

Noyes, P. D., Hinton, D. E., & Stapleton, H. M. (2011). Accumulation and Debromination of Decabromodiphenyl Ether (BDE-209) in Juvenile Fathead Minnows (Pimephales promelas) Induces Thyroid Disruption and Liver Alterations. Toxicological Sciences, 122(2), 265–274. 10.1093/toxsci/kfr105

Oaks, J. L., Gilbert, M., Virani, M. Z., Watson, R. T., Meteyer, C. U., Rideout, B. A., Shivaprasad, H. L., Ahmed, S., Iqbal Chaudhry, M. J., Arshad, M., Mahmood, S., Ali, A., & Ahmed Khan, A. (2004). Diclofenac residues as the cause of vulture population decline in Pakistan. Nature, 427(6975), 630–633. 10.1038/nature02317

Parrott, J. L., Bartlett, A. J., & Balakrishnan, V. K. (2016). Chronic toxicity of azo and anthracenedione dyes to embryo-larval fathead minnow. Environmental Pollution, 210, 40–47. 10.1016/j.envpol.2015.11.037

Pastorino, P., Prearo, M., & Barceló, D. (2024). Ethical principles and scientific advancements: In vitro, in silico, and non-vertebrate animal approaches for a green ecotoxicology. Green Analytical Chemistry, 8, 100096. 10.1016/j.greeac.2024.100096

Patterson, E. A., Whelan, M. P., & Worth, A. P. (2021a). The role of validation in establishing the scientific credibility of predictive toxicology approaches intended for regulatory application. Computational Toxicology, 17, 100144. 10.1016/j.comtox.2020.100144

Patterson, E. A., Whelan, M. P., & Worth, A. P. (2021b). The role of validation in establishing the scientific credibility of predictive toxicology approaches intended for regulatory application. Computational Toxicology, 17, 100144. 10.1016/j.comtox.2020.100144

Persson, L., Carney Almroth, B. M., Collins, C. D., Cornell, S., de Wit, C. A., Diamond, M. L., Fantke, P., Hassellöv, M., MacLeod, M., Ryberg, M. W., Søgaard Jørgensen, P., Villarrubia-Gómez, P., Wang, Z., & Hauschild, M. Z. (2022). Outside the Safe Operating Space of the Planetary Boundary for Novel Entities. Environmental Science & Technology, 56(3), 1510–1521. 10.1021/acs.est.1c04158

Posthuma, L., Brack, W., van Gils, J., Focks, A., Müller, C., de Zwart, D., & Birk, S. (2019). Mixtures of chemicals are important drivers of impacts on ecological status in European surface waters. Environmental Sciences Europe, 31(1), 71. 10.1186/s12302-019-0247-4

Prakash, R. (2015). Chapter 1 - General Aspects of Organofluorine Compounds. In V. P. Reddy (Ed.), Organofluorine Compounds in Biology and Medicine (pp. 1–27). Elsevier. 10.1016/B978-0-444-53748-5.00001-0

Price, P. S., Hubbell, B. J., Hagiwara, S., Paoli, G. M., Krewski, D., Guiseppi-Elie, A., Gwinn, M. R., Adkins, N. L., & Thomas, R. S. (2022). A Framework that Considers the Impacts of Time, Cost, and Uncertainty in the Determination of the Cost Effectiveness of Toxicity-Testing Methodologies. Risk Analysis, 42(4), 707–729. 10.1111/risa.13810

Probst, D., & Reymond, J.-L. (2018). A probabilistic molecular fingerprint for big data settings. Journal of Cheminformatics, 10(1), 66. 10.1186/s13321-018-0321-8

Probst, D., & Reymond, J.-L. (2020). Visualization of very large high-dimensional data sets as minimum spanning trees. Journal of Cheminformatics, 12(1), 12. 10.1186/s13321-020-0416-x

REACH Regulation. (2025). Consolidated text: Regulation (EC) No 1907/2006 of the European Parliament and of the Council of 18 December 2006 concerning the Registration, Evaluation, Authorisation and Restriction of Chemicals (REACH), establishing a European Chemicals Agency, amending Directive 1999/45/EC and repealing Council Regulation (EEC) No 793/93 and Commission Regulation (EC) No 1488/94 as well as Council Directive 76/769/EEC and Commission Directives 91/155/EEC, 93/67/EEC, 93/105/EC and 2000/21/EC.

Rutz, A., Sorokina, M., Galgonek, J., Mietchen, D., Willighagen, E., Gaudry, A., Graham, J. G., Stephan, R., Page, R., Vondrášek, J., Steinbeck, C., Pauli, G. F., Wolfender, J.-L., Bisson, J., & Allard, P.-M. (2022). The LOTUS initiative for open knowledge management in natural products research. ELife, 11, e70780. 10.7554/eLife.70780

Sigmund, G., Ågerstrand, M., Antonelli, A., Backhaus, T., Brodin, T., Diamond, M. L., Erdelen, W. R., Evers, D. C., Hofmann, T., Hueffer, T., Lai, A., Torres, J. P. M., Mueller, L., Perrigo, A. L., Rillig, M. C., Schaeffer, A., Scheringer, M., Schirmer, K., Tlili, A., … Groh, K. J. (2023). Addressing chemical pollution in biodiversity research. Global Change Biology, 29(12), 3240–3255. 10.1111/gcb.16689

Thornton, L. M., Path, E. M., Nystrom, G. S., Venables, B. J., & Sellin Jeffries, M. K. (2018). Embryo-larval BDE-47 exposure causes decreased pathogen resistance in adult male fathead minnows (Pimephales promelas). Fish & Shellfish Immunology, 80, 80–87. 10.1016/j.fsi.2018.05.059

Tsai, Y.-H., & Lein, P. J. (2021). Mechanisms of organophosphate neurotoxicity. Current Opinion in Toxicology, 26, 49–60. 10.1016/j.cotox.2021.04.002

Tukaj, Z., & Aksmann, A. (2007). Toxic effects of anthraquinone and phenanthrenequinone upon Scenedesmus strains (green algae) at low and elevated concentration of CO2. Chemosphere, 66(3), 480–487. 10.1016/j.chemosphere.2006.05.072

van de Meent, D., de Zwart, D., & Posthuma, L. (2020). Screening-Level Estimates of Environmental Release Rates, Predicted Exposures, and Toxic Pressures of Currently Used Chemicals. Environmental Toxicology and Chemistry, 39(9), 1839–1851. 10.1002/etc.4801

van der Veen, I., & de Boer, J. (2012). Phosphorus flame retardants: Properties, production, environmental occurrence, toxicity and analysis. Chemosphere, 88(10), 1119–1153. 10.1016/j.chemosphere.2012.03.067

Wang, Z., Walker, G. W., Muir, D. C. G., & Nagatani-Yoshida, K. (2020). Toward a Global Understanding of Chemical Pollution: A First Comprehensive Analysis of National and Regional Chemical Inventories. Environmental Science & Technology, 54(5), 2575–2584. 10.1021/acs.est.9b06379

Williams, A. J., Grulke, C. M., Edwards, J., McEachran, A. D., Mansouri, K., Baker, N. C., Patlewicz, G., Shah, I., Wambaugh, J. F., Judson, R. S., & Richard, A. M. (2017). The CompTox Chemistry Dashboard: a community data resource for environmental chemistry. Journal of Cheminformatics, 9(1), 61. 10.1186/s13321-017-0247-6

Wittwehr, C., Blomstedt, P., Gosling, J. P., Peltola, T., Raffael, B., Richarz, A.-N., Sienkiewicz, M., Whaley, P., Worth, A., & Whelan, M. (2020). Artificial Intelligence for chemical risk assessment. Computational Toxicology, 13, 100114. 10.1016/j.comtox.2019.100114

Yang, H.-B., Zhao, Y.-Z., Tang, Y., Gong, H.-Q., Guo, F., Sun, W.-H., Liu, S.-S., Tan, H., & Chen, F. (2019). Antioxidant defence system is responsible for the toxicological interactions of mixtures: A case study on PFOS and PFOA in Daphnia magna. Science of The Total Environment, 667, 435–443. 10.1016/j.scitotenv.2019.02.418

Yang, J., Zhao, Y., Li, M., Du, M., Li, X., & Li, Y. (2019). A review of a class of emerging contaminants: the classification, distribution, intensity of consumption, synthesis routes, environmental effects and expectation of pollution abatement to organophosphate flame retardants (OPFRs). International Journal of Molecular Sciences, 20(12), 2874.

Yang, Li, Y.-X., Ren, F.-Z., Luo, J., & Pang, G.-F. (2019). Assessment of the endocrine-disrupting effects of organophosphorus pesticide triazophos and its metabolites on endocrine hormones biosynthesis, transport and receptor binding in silico. Food and Chemical Toxicology, 133, 110759. 10.1016/j.fct.2019.110759

Zhang, Q., Yang, L., Wang, H., Wu, C., Cao, R., Zhao, M., Su, G., & Wang, C. (2024). A comprehensive evaluation of the endocrine-disrupting effects of emerging organophosphate esters. Environment International, 193, 109120. 10.1016/j.envint.2024.109120

Zhou, J., Yun, X., Wang, J., Li, Q., & Wang, Y. (2022). A review on the ecotoxicological effect of sulphonamides on aquatic organisms. Toxicology Reports, 9, 534–540. 10.1016/j.toxrep.2022.03.034

