## Supplemental Materials for "A chemical space model for the exploration of eco-toxicological data"

### Supplementary Materials

Lopez-Rodriguez D[1,2], Guerrero-Limón G[3], Chèvre N[1]

[1] Ecotoxicology Group, Institute of Earth Surface Dynamics, Faculty of Geosciences and the Environment, University of Lausanne, 1015 Lausanne, Switzerland

[2] Department of Biomedical Science, University of Lausanne, Lausanne, Switzerland

[3] Vitis Regulatory, Brussels, Belgium

**Supplementary Materials**

**Figure S1**. Chemical space exploration of physicochemical properties. Representation of the number of aliphatic carbocycles (**A**), minimum electrotopological state index (**B**), monoisotopic mass (**C**), number of heavy atoms (**D**), number of heteroatoms (**E**), BalabanJ index (**F**) and Bertz chemical complexity index (**G**).

**Figure S2**. Organofluoride and organoiodide toxicity in *Daphnia magna*. TMAP displaying organofluoride toxicity distribution in *Daphnia magna* (**A**) and chemical classification within the chemical subspace (**B**). Top organoiodide (**C**) and pyrethroid (**D**) substances predicted to be toxic for *Daphnia magna*.

**Figure S3**. Anthraquinone and benzoic acid ester toxicity in *Pimephales Promelas*. TMAP displaying anthraquinone toxicity distribution in *Pimephales Promelas* (**A**) and chemical classification within the chemical subspace (**B**). Top anthraquinone (**C**) substances predicted to be toxic for *Pimephales Promelas*. (**D-E**) TMAP displaying benzoic acid ester toxicity distribution in *Pimephales Promelas* and chemical classification within the chemical subspace. (**F**) Most toxic benzoic acid ester substances predicted to be toxic for *Pimephales Promelas*.

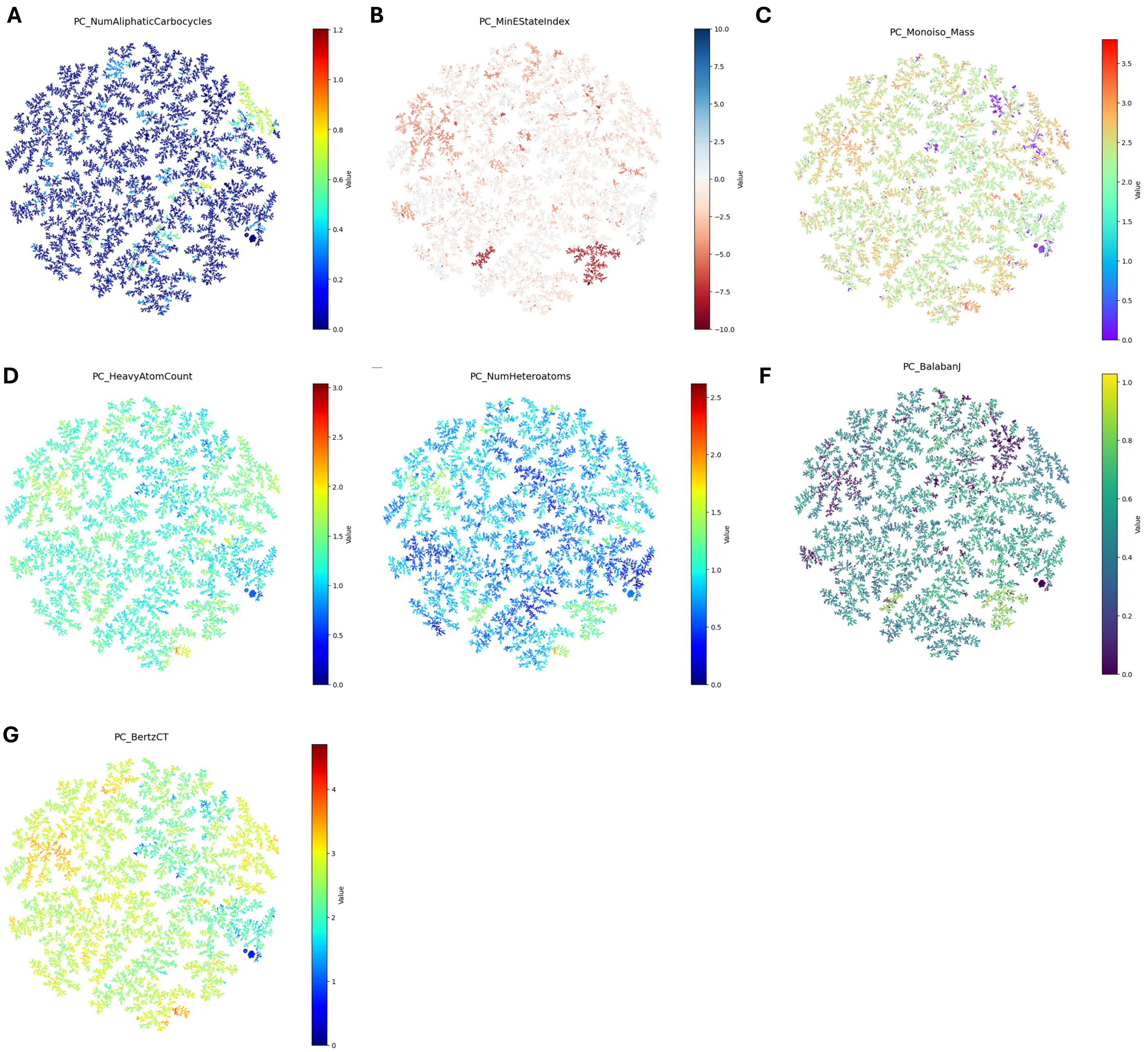
A
PC_NumAliphaticCarbocycles
B
PC_MinEStateIndex
C
PC_Monoiso_Mass
D
PC_HeavyAtomCount
PC_NumHeteroatoms
F
PC_BalabanJ
G
PC_BertzCT
Value

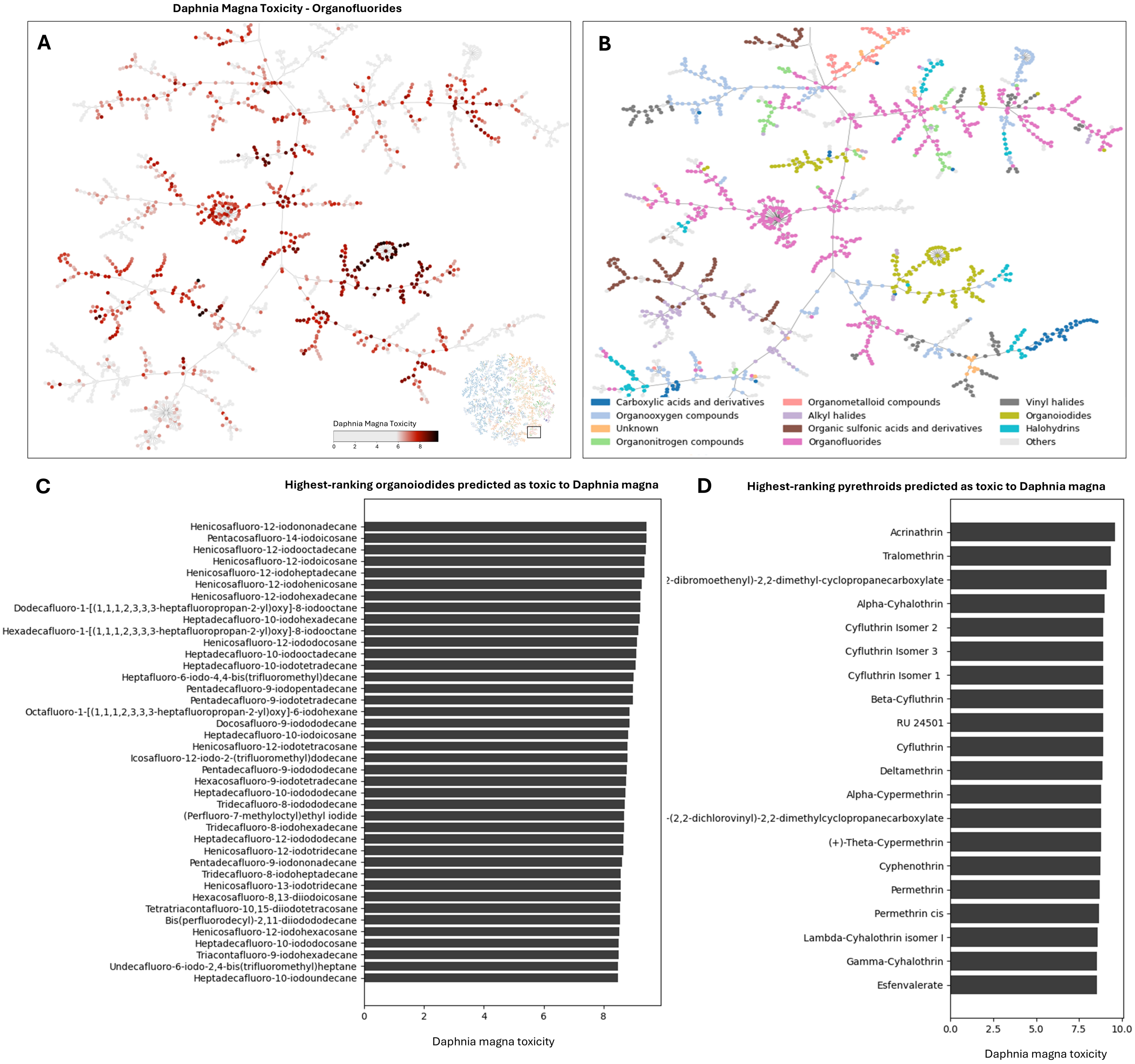
Daphnia Magna Toxicity - Organofluorides
A
Daphnia Magna Toxicity
0 2 4 6 8
B
Carboxylic acids and derivatives
Organooxygen compounds
Unknown
Organonitrogen compounds
Organometalloid compounds
Alkyl halides
Organic sulfonic acids and derivatives
Organofluorides
Vinyl halides
Organoiodides
Halohydrins
Others
C
Highest-ranking organoiodides predicted as toxic to Daphnia magna
Henicosafluoro-12-iodononadecane
Pentacosafluoro-14-iodoicosane
Henicosafluoro-12-iodooctadecane
Henicosafluoro-12-iodoicosane
Henicosafluoro-12-iodoheptadecane
Henicosafluoro-12-iodohenicosane
Henicosafluoro-12-iodohexadecane
Dodecafluoro-1-[(1,1,1,2,3,3,3-heptafluoropropan-2-yl)oxy]-8-iodooctane
Heptadecafluoro-10-iodohexadecane
Hexadecafluoro-1-[(1,1,1,2,3,3,3-heptafluoropropan-2-yl)oxy]-8-iodooctane
Henicosafluoro-12-iodododocosane
Heptadecafluoro-10-iodooctadecane
Heptadecafluoro-10-iodotetradecane
Heptafluoro-6-iodo-4,4-bis(trifluoromethyl)decane
Pentadecafluoro-9-iodopentadecane
Pentadecafluoro-9-iodotetradecane
Octafluoro-1-[(1,1,1,2,3,3,3-heptafluoropropan-2-yl)oxy]-6-iodohexane
Docosafluoro-9-iodododecane
Heptadecafluoro-10-iodoicosane
Henicosafluoro-12-iodotetracosane
Icosafluoro-12-iodo-2-(trifluoromethyl)dodecane
Pentadecafluoro-9-iodododecane
Hexacosafluoro-9-iodotetradecane
Heptadecafluoro-10-iodododecane
Tridecafluoro-8-iodododecane
(Perfluoro-7-methyloctyl)ethyl iodide
Tridecafluoro-8-iodohexadecane
Heptadecafluoro-12-iodododecane
Henicosafluoro-12-iodotridecane
Pentadecafluoro-9-iodononadecane
Tridecafluoro-8-iodoheptadecane
Henicosafluoro-13-iodotridecane
Hexacosafluoro-8,13-diiodoicosane
Tetratriacontafluoro-10,15-diiodotetracosane
Bis(perfluorodecyl)-2,11-diiodododecane
Henicosafluoro-12-iodohexacosane
Heptadecafluoro-10-iodododocosane
Triacontafluoro-9-iodohexadecane
Undecafluoro-6-iodo-2,4-bis(trifluoromethyl)heptane
Heptadecafluoro-10-iodoundecane
0 2 4 6 8
Daphnia magna toxicity
D
Highest-ranking pyrethroids predicted as toxic to Daphnia magna
Acrinathrin
Tralomethrin
?-dibromoethenyl)-2,2-dimethyl-cyclopropanecarboxylate
Alpha-Cyhalothrin
Cyfluthrin Isomer 2
Cyfluthrin Isomer 3
Cyfluthrin Isomer 1
Beta-Cyfluthrin
RU 24501
Cyfluthrin
Deltamethrin
Alpha-Cypermethrin
-(2,2-dichlorovinyl)-2,2-dimethylcyclopropanecarboxylate
(+)-Theta-Cypermethrin
Cyphenothrin
Permethrin
Permethrin cis
Lambda-Cyhalothrin isomer I
Gamma-Cyhalothrin
Esfenvalerate
0.0 2.5 5.0 7.5 10.0
Daphnia magna toxicity

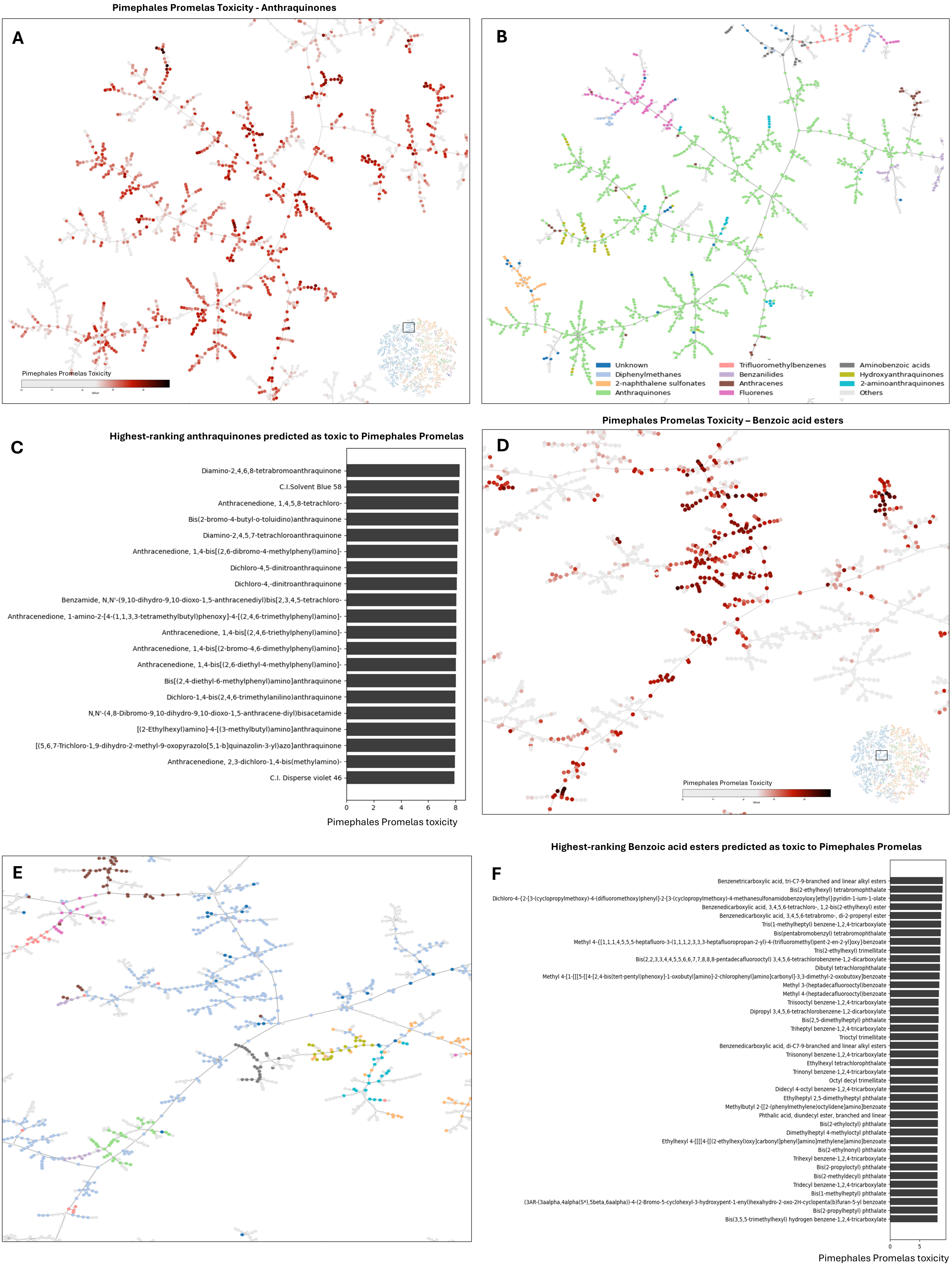

Pimephales Promelas Toxicity - Anthraquinones
A
Pimephales Promelas Toxicity
B
Unknown
Diphenylmethanes
2-naphthalene sulfonates
Anthraquinones
Trifluoromethylbenzenes
Benzanilides
Anthracenes
Fluorenes
Aminobenzoic acids
Hydroxyanthraquinones
2-aminoanthraquinones
Others
Highest-ranking anthraquinones predicted as toxic to Pimephales Promelas
C
Diamino-2,4,6,8-tetrabromoanthraquinone
C.I.Solvent Blue 58
Anthracenedione, 1,4,5,8-tetrachloro-
Bis(2-bromo-4-butyl-o-toluidino)anthraquinone
Diamino-2,4,5,7-tetrachloroanthraquinone
Anthracenedione, 1,4-bis[(2,6-dibromo-4-methylphenyl)amino]-
Dichloro-4,5-dinitroanthraquinone
Dichloro-4,-dinitroanthraquinone
Benzamide, N,N'-(9,10-dihydro-9,10-dioxo-1,5-anthracenediyl)bis[2,3,4,5-tetrachloro-
Anthracenedione, 1-amino-2-[4-(1,1,3,3-tetramethylbutyl)phenoxy]-4-[(2,4,6-trimethylphenyl)amino]-
Anthracenedione, 1,4-bis[(2,4,6-triethylphenyl)amino]-
Anthracenedione, 1,4-bis[(2-bromo-4,6-dimethylphenyl)amino]-
Anthracenedione, 1,4-bis[(2,6-diethyl-4-methylphenyl)amino]-
Bis[(2,4-diethyl-6-methylphenyl)amino]anthraquinone
Dichloro-1,4-bis(2,4,6-trimethylanilino)anthraquinone
N,N'-(4,8-Dibromo-9,10-dihydro-9,10-dioxo-1,5-anthracene-diyl)bisacetamide
[(2-Ethylhexyl)amino]-4-[(3-methylbutyl)amino]anthraquinone
[(5,6,7-Trichloro-1,9-dihydro-2-methyl-9-oxopyrazolo[5,1-b]quinazolin-3-yl)azo]anthraquinone
Anthracenedione, 2,3-dichloro-1,4-bis(methylamino)-
C.I. Disperse violet 46
0
2
4
6
8
Pimephales Promelas toxicity
Pimephales Promelas Toxicity – Benzoic acid esters
D
Pimephales Promelas Toxicity
E
Highest-ranking Benzoic acid esters predicted as toxic to Pimephales Promelas
F
Benzenetricarboxylic acid, tri-C7-9-branched and linear alkyl esters
Bis(2-ethylhexyl) tetrabromophthalate
Dichloro-4-{2-[3-(cyclopropylmethoxy)-4-(difluoromethoxy)phenyl]-2-[3-(cyclopropylmethoxy)-4-methanesulfonamidobenzoyloxy]ethyl}pyridin-1-ium-1-olate
Benzenedicarboxylic acid, 3,4,5,6-tetrachloro-, 1,2-bis(2-ethylhexyl) ester
Benzenedicarboxylic acid, 3,4,5,6-tetrabromo-, di-2-propenyl ester
Tris(1-methylheptyl) benzene-1,2,4-tricarboxylate
Bis(pentabromobenzyl) tetrabromophthalate
Methyl 4-{[1,1,1,4,5,5,5-heptafluoro-3-(1,1,1,2,3,3,3-heptafluoropropan-2-yl)-4-(trifluoromethyl)pent-2-en-2-yl]oxy}benzoate
Tris(2-ethylhexyl) trimellitate
Bis(2,2,3,3,4,4,5,5,6,6,7,7,8,8,8-pentadecafluorooctyl) 3,4,5,6-tetrachlorobenzene-1,2-dicarboxylate
Dibutyl tetrachlorophthalate
Methyl 4-[1-[[[5-[[4-[2,4-bis(tert-pentyl)phenoxy]-1-oxobutyl]amino]-2-chlorophenyl]amino]carbonyl]-3,3-dimethyl-2-oxobutoxy]benzoate
Methyl 3-(heptadecafluorooctyl)benzoate
Methyl 4-(heptadecafluorooctyl)benzoate
Triisooctyl benzene-1,2,4-tricarboxylate
Dipropyl 3,4,5,6-tetrachlorobenzene-1,2-dicarboxylate
Bis(2,5-dimethylheptyl) phthalate
Triheptyl benzene-1,2,4-tricarboxylate
Trioctyl trimellitate
Benzenedicarboxylic acid, di-C7-9-branched and linear alkyl esters
Triisononyl benzene-1,2,4-tricarboxylate
Ethylhexyl tetrachlorophthalate
Trinonyl benzene-1,2,4-tricarboxylate
Octyl decyl trimellitate
Didecyl 4-octyl benzene-1,2,4-tricarboxylate
Ethylheptyl 2,5-dimethylheptyl phthalate
Methylbutyl 2-[[2-(phenylmethylene)octylidene]amino]benzoate
Phthalic acid, diundecyl ester, branched and linear
Bis(2-ethyloctyl) phthalate
Dimethylheptyl 4-methyloctyl phthalate
Ethylhexyl 4-[[[[4-[[(2-ethylhexyl)oxy]carbonyl]phenyl]amino]methylene]amino]benzoate
Bis(2-ethylnonyl) phthalate
Trihexyl benzene-1,2,4-tricarboxylate
Bis(2-propyloctyl) phthalate
Bis(2-methyldecyl) phthalate
Tridecyl benzene-1,2,4-tricarboxylate
Bis(1-methylheptyl) phthalate
(3AR-(3aalpha,4alpha(S*),5beta,6aalpha))-4-(2-Bromo-5-cyclohexyl-3-hydroxypent-1-enyl)hexahydro-2-oxo-2H-cyclopenta(b)furan-5-yl benzoate
Bis(2-propylheptyl) phthalate
Bis(3,5,5-trimethylhexyl) hydrogen benzene-1,2,4-tricarboxylate
0
5
Pimephales Promelas toxicity
